# Development of an oral swab based microbiome test for the detection of feline dental disease

**DOI:** 10.1101/2021.04.23.441192

**Authors:** Damian Kao, Julie Yang, Sean Nisperos, Norma Drew, Polina Berezovskaya, Kaushalya Kuruppu, Yuliana Mihaylova

**Affiliations:** Basepaws, 1124 W Carson Street, MRL Building, 3rd floor, Torrance, CA 90502

**Keywords:** oral microbiome, feline dental disease, tooth resorption, periodontal disease, halitosis

## Abstract

Variations in the microbial composition of the mouth (the oral microbiome) have known associations with dental and systemic disease. While this is relatively well understood in humans, research on this topic in companion animals, and in cats in particular, has been limited. In this study, we used oral microbiome data obtained from shotgun metagenomic sequencing of 38,000 cats (data gathered through a direct-to-consumer cat DNA testing platform) to reveal the staggering diversity of the feline oral microbiome, identifying 8,344 microbial species across the entire cohort. We used a subset of these data points (6,110 cats) to develop a feline dental health test able to assess a cat’s risk of having periodontal disease, tooth resorption and halitosis based on their oral microbiome. After filtering out classified microbial reads with low abundance, we were able to detect 606 microbes in a single cat’s oral microbiome, identifying not just bacteria, but also viruses, fungi, archaea and protozoa. Due to the shortage of available published research on the microbial signature of tooth resorption and halitosis in cats, we used our periodontal disease feline cohort (n=570) to validate our approach. We observed microbial compositional abundance trends consistent with previously reported findings from feline, canine and human studies on periodontal disease. We used compositional abundance-based statistical methods relying on pairwise log-ratio (PLR) transformation to identify microbes significantly correlated with each of the three dental conditions of interest. We identified a set of 27 microbes that are predictive for all three dental conditions, as well as microbes specifically predictive of periodontal disease, tooth resorption or halitosis. We used the compositional abundance profiles of predictive microbes to develop a risk score based model assessing the probability that a cat is suffering from each of the three dental conditions. The model had highest sensitivity for halitosis (72%) and highest specificity for tooth resorption (78%). Lastly, we observed relatively consistent dental disease risk profiles when we compared data from sample collection methods targeting the whole mouth versus those targeting the gum line specifically. In contrast, samples collected in triplicates from the same cats using a sampling method targeting the whole mouth showed more variation in the generated risk profiles. This was likely due to a failure to consistently collect sufficient sample material from areas of the mouth where microbes relevant to dental pathology would be found in highest amounts (i.e., the gum line). For this reason, we have modified the test’s instructions to emphasize the importance of targeting the gum line during sample collection. Regular at home or in clinic screening with the feline dental health test described in this study has the potential to facilitate early detection and prevention of dental disease.

## Introduction

Nutritional and environmental factors, as well as disease states, play an important role in the dynamic microbial composition of a cat’s mouth (their oral microbiome). With the mouth being the first line of defense from a constant exposure to foreign microbes, its microbiome has evolved to be competitive and territorial^1^. It is comprised of microbes that excel at defending their territory and are typically able to avoid being replaced by foreign invaders, including pathogens. However, dysbiosis inducing events such as poor diet or poor dental hygiene, can lead to pathogenic microbes colonizing disproportionately large parts of the oral cavity, which can be associated with pathology. Understanding the composition of the oral microbiome can provide information about the health of oral tissues and point to potential dental and gum diseases^2^. It is now recognised that most dental diseases are associated with complex interactions involving a multitude of microbes, as opposed to a single microbe^3^.

The field of oral microbiome research in companion animals has received little focus and it is still in its infancy, as a result. Existing studies base their conclusions on small sample sizes^3–6^ and outdated culture-based techniques for querying the microbiome^4, 7–9^. It is estimated that only around 2% of all existing bacteria can be cultured in the laboratory^10^, meaning that in studies relying on this method for microbial classification, many important microbial organisms will likely be missed, while false emphasis might be placed on particular species, simply because they could be cultured. This problem is compounded by the fact that lab culturing provides a very bacteria-centric view of the microbiome, often ignoring other microorganisms such as fungi, protozoa, archaea and viruses^11^. To alleviate some of these issues, a non-culture dependent technique called 16S rRNA gene sequencing, which relies on Next Generation Sequencing (NGS), was introduced to study microbial populations. While the technique only allows classification of bacteria and archaea (often only to the genus level^12^), it is a significant improvement over bacterial culturing studies of the microbiome. However, the gold standard for the comprehensive study of the microbiome is shotgun metagenomic sequencing^13, 14^, which allows capturing complete or near-complete genomes of organisms across all domains of life, not just bacteria and archaea. This method also allows microbial classification down to the species or, in some instances, strain level, unlike 16S gene sequencing.

Using shotgun metagenomic oral microbiome sequencing of 38,000 domestic cats and compositional data analysis techniques, we performed a comprehensive survey of the feline oral microbiome, identifying 8,344 microbial species. We used these data in conjunction with user provided health history information on the cats to develop a feline dental health test able to assess a cat’s likelihood of suffering from periodontal disease, tooth resorption or halitosis based on the microbial profile of their mouth. The test relies on a painless oral swab sample collection, does not require anesthetizing the animal, and can be performed by the pet owner at their home or by the veterinarian at the clinic. In line with what studies in humans have shown when the oral microbiome is surveyed via buccal, supragingival or subgingival sample collection methods, our feline oral microbiome test could serve as an early indicator of dental disease-associated processes not yet visible to the naked eye^15^. Routine use of our feline dental health test has the potential to facilitate diagnosis of early stage dental diseases, driving more cats to the veterinary office early on and reducing the number of emergency dental vet visits in the long run.

## Materials and Methods

### Ethics statement

The present study is of observational nature and does not utilize any invasive procedures. All feline oral swab samples and accompanying health history information used in this study were provided voluntarily by pet owners who agreed in electronic form for their cat’s data to be used in an aggregated de-identified format for research purposes.

### Domestic cat cohort used for oral microbiome database construction

The study cohort consisted of de-identified domestic cats whose owners had purchased an oral swab-based cat DNA test. Our initial sample size was 38,000 cats. For data consistency purposes, we selected only those samples processed with a ligation-based library preparation method (n=15,154) for further analysis and excluded those processed with a tagmentation-based library preparation method. This was done due to an observed effect of the library preparation method on microbial species richness (**Supplementary figure 1**). The ligation-based method was preferred because the number of sequencing reads per sample had minimal impact on the number of microbial species detected. In addition, Tn5 transposase assisted tagmentation is known to introduce GC sequencing bias, particularly in metagenomic communities^16^. Of the remaining feline samples prepared with a ligation-based library prep, 6,962 had matching user-provided health and traits (phenotype) data. The phenotype data was provided by the pet owner in the form of an online questionnaire. For this study, we sub-selected cats whose owners provided details about their pet’s dental health history. We focused on the following cohorts:

- **PD cohort:** Cats reported by their owners to have been diagnosed by a veterinarian with **p**eriodontal **d**isease (n=570, after all data filtering steps described in the *Data analysis* section were performed)
- **TR cohort:** Cats reported by their owners to have been diagnosed by a veterinarian with **t**ooth **r**esorption (n=111, after all data filtering steps described in the *Data analysis* section were performed)
- **BB cohort:** Cats reported by their owners to have **b**ad **b**reath (n=137, after all data filtering steps described in the *Data analysis* section were performed), also characterized specifically as ‘death and decay’ breath
- **Healthy cohort:** Cats 1-3 years of age, with no diagnosed dental or general health conditions (n=1,147, after all data filtering steps described in the *Data analysis* section were performed). Cats below one year of age were intentionally excluded from this group with the purpose of avoiding any potential kitten-specific oral microbiome bias. Since age is a known predictive factor for dental and general diseases, the 1-3 age range was selected to minimize the possibility that older cats with yet undiagnosed diseases could be misclassified as healthy cats.
- **TB cohort:** Cats reported by their owners to have ‘**t**ypical’ cat **b**reath (n=4,109, after all data filtering steps described in the *Data analysis* section were performed)

### Domestic cat cohorts used for Study 1 and Study 2

Twelve cats took part in Study 1 and eleven in Study 2. Participants were recruited through an email inviting participation in studies focused on feline dental health. Participants in Study 1 received 2 DNAGenotek’s PERFORMAgene (PG-100) oral swab collection devices and were instructed to swab their cat once using each swab. The first swab was used to collect a sample from the whole mouth, while the second one targeted the gum line specifically. Participants were also asked to collect the two samples in one sitting. Participants in Study 2 received 3 DNAGenotek’s PERFORMAgene (PG-100) oral swab collection devices and were instructed to perform three whole mouth oral swab collections in one sitting.

### Oral swab sample collection

The vast majority of feline oral swab samples were collected by pet owners at their respective homes, with a small proportion of sample collections performed by a veterinarian. Pet owners and veterinarians were instructed to collect the samples at least 30 minutes to an hour after the cat had had anything to eat or drink. They were also instructed to keep the oral swab sample collection device in the cat’s mouth for at least 5 seconds. The sample collection device used was DNAGenotek’s PERFORMAgene (PG-100).

### DNA extraction

Metagenomic DNA was extracted from feline oral samples via heat activation (55°C) for an hour on a shaker, followed by SPRI magnetic beads-based DNA extraction (MCLAB, MBC-200) using 80% ethanol for purification. The DNA was quantified using a GloMax Plate Reader (Promega).

### Next Generation Sequencing (NGS) library preparation

Following metagenomic DNA extraction and quantification, each sample was prepared for NGS using the LOTUS DNA library prep kit (IDT) following the manufacturer’s instructions. Each sample was dual-barcoded with iTRU indices^17^. The prepared sequencing libraries were quantified using a GloMax Plate Reader (Promega) and equal-mass pooled into 96-sample pools. The pools were then visualized (to assess fragment size distribution) and quantified using a 2100 Bioanalyzer instrument (Agilent).

### NGS

Following standard QC steps, the 96-sample pools were loaded onto an Illumina HiSeq X or NovaSeq 6000 Next Generation Sequencing machine, as per the manufacturer’s protocols.

### Data analysis - sequencing read classification and data filtering

The raw sequencing data was demultiplexed and trimmed to remove low-quality data using the program Trimmomatic 0.32^18^. The data was then mapped to the latest version of the feline genome Felis_catus_9.0^19^. For every sample, there were 5-7% sequencing reads that did not map to the feline genome. We classified these unmapped reads using the KRAKEN2 metagenomic sequence classifier to identify the microbial organisms present in each sample^20^. The confidence score of was used as a cutoff for the KRAKEN2 classification algorithm, as recommended by the KRAKEN2 algorithm’s authors. We filtered out all samples with fewer than 10,000 classified microbial reads or more than 500,000 classified microbial reads. We also filtered out the reads for microbial species with a non-zero mean of fewer than 10 reads. We then used Bracken^21^, a statistical method for calculating species abundance in DNA sequencing data from a metagenomic sample.

### Data analysis - dental disease risk prediction and species compositional abundance assessment

To assess over- or under-representation of particular microbial species in periodontal disease, the Bracken output data underwent Centered Log-Ratio (CLR) transformation. This was done to account for potential compositional biases, which are a well established problem in microbiome data analysis^22^. A z-test was then performed on the CLR transformed data to identify microbial species with statistically significant increased and decreased compositional abundance in periodontal disease compared to control.

As a first step towards identifying microbes significantly correlated with each dental condition, Pairwise Log-Ratio (PLR) transformation^23^ was performed on the Bracken output species level read counts. Next, we identified the significant PLR comparisons (p-value < 0.01) between control and condition by performing a z-test. The following comparisons between condition and control were performed:

- PD cohort vs Healthy cohort
- TR cohort vs Healthy cohort
- BB cohort vs TB cohort

We assessed the frequency of each microbial species in all significant PLRs. We kept only microbial species where 50% or more of their maximum possible comparisons with other species were significant. This measure was used as a proxy for the importance of different microbial species in the three dental conditions of interest. We called these microbial species ‘predictive’ for each respective dental condition. In order to identify population-wide microbial compositional abundance patterns characteristic of periodontal disease, tooth resorption or halitosis, for each of the three conditions, we scored each sample by comparing the predictive pairwise log-ratios (pPLRs) of the sample to the mean pPLRs of controls, taking into account the direction and magnitude of the difference. Next, we fitted 3 Gaussian mixture models (one for each dental condition) with 2 components each - healthy cohort (or TB cohort) and dental condition - onto the distribution of the average log ratio difference score between pairwise microbial interactions. This modeling approach generates a 0 to 1 score for each sample, which represents the probability that the sample belongs to the control cohort or to the respective dental condition cohort. This is the model we used to assess a cat’s risk of having tooth resorption, periodontal disease or halitosis based on the pattern of predictive microbe PLRs observed in their oral microbiome. We applied the following 3 risk assessment categories based on the probability score generated for each sample: the 0.0 - 0.33 bracket is classified as ‘low risk’ of having a dental condition, >0.33 - 0.66 is classified as ‘medium risk’ and >0.66 - 1.0 is classified as ‘high risk’.

## Results

### Developing a feline oral microbiome test and constructing a reference database

We developed a painless oral swab-based feline dental health test relying on metagenomic shotgun NGS (**Figure 1A**). The test compares a cat’s oral microbiome to the oral microbiomes of cats reported by their owners to have been diagnosed with tooth resorption or periodontal disease, or to have bad breath characterized by a ‘death and decay’ odour. To make this comparison possible, we built a reference database of feline oral microbiomes (**Figure 1B**). We gathered oral microbiome data from 38,000 domestic cats, which allowed us to identify 8,344 microbial species that can be found in a cat’s mouth. On average, we identified 606 microbial species per cat, 97.% of which were classified as bacteria and archaea, 0.27% as DNA viruses (RNA viruses cannot be detected with shotgun metagenomic sequencing), 0.02% as phages and <2% as fungi.

**Figure 1:**
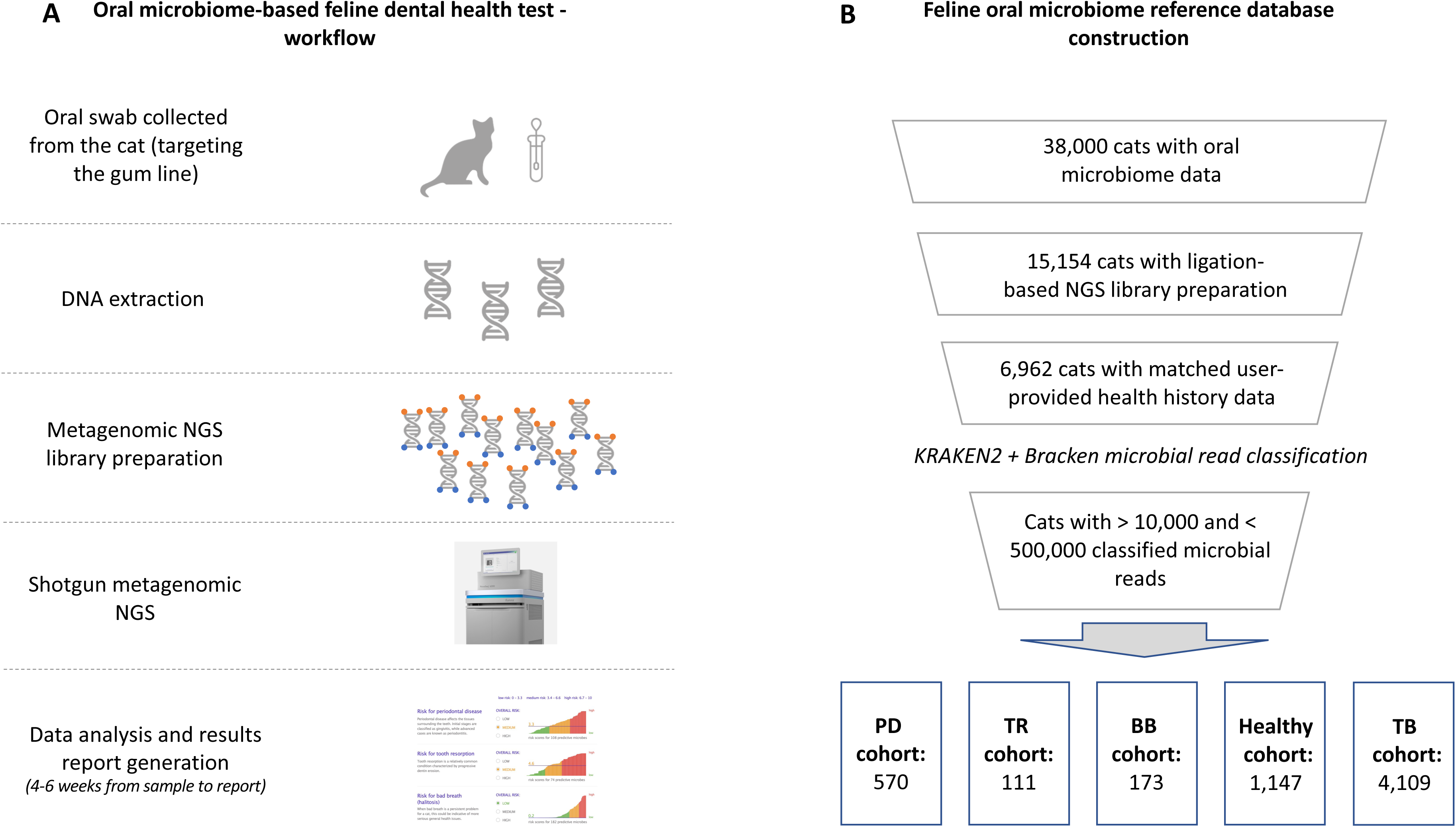
Feline dental health test workflow and oral microbiome reference database construction. (A) The feline dental health test workflow includes collecting an oral swab from the cat in a DNA preservation solution, extracting and preparing the DNA for shotgun metagenomic NGS, sequencing, data analysis and report generation. (B) The feline oral microbiome reference database was constructed through applying sequential filters on our initial database of 38,000 cats. First, we removed all data from tagmentation-based NGS library preparation samples, then we excluded samples where we did not have an accompanying phenotype/health history record for the cat. After we classified the microbial reads in each sample using KRAKEN2 and Bracken, we removed samples with fewer than 10,000 and more than 500,000 classified microbial reads. This resulted in a periodontal disease (PD) cohort of 570 cats, tooth resorption (TR) cohort of 111 cats, bad breath (BB) cohort of 173 cats, healthy cohort of 1,147 cats and typical breath (TB) cohort of 4,109 cats.

After we performed the filtering steps described in the **Materials and Methods** section and **Figure 1B**, we were left with 6,110 cats in our final dental health oral microbiome database. The cats were split into those diagnosed with periodontal disease and no known additional dental diseases (PD cohort), those diagnosed with tooth resorption and no known additional dental diseases (TR cohort), those reported to have bad breath by their pet owners (BB cohort), and those reported to have typical cat breath by their owners (TB cohort).

### Replicating and further characterizing the microbial signature of feline periodontal disease reported in literature

Since our database relied on pet owner reported data and not on diagnoses provided directly from the veterinarian, we wanted to validate our findings by comparing the dental disease related microbial signatures we observed in our data to those reported in literature. Unfortunately, there is no available literature describing the state of the feline oral microbiome when a diagnosis of tooth resorption or halitosis is present. However, there is already existing knowledge regarding the key microbial compositional abundance trends characteristic of periodontal disease in cats, dogs and humans^2–5, 7^. Some of these trends include increased abundance of bacterial species *Porphyromonas gingivalis*^4, 7–9, 24, 25^, *Tannerella forsythia*^25^*, Bacteroides zoogleoformans*^26, 27^*, Desulfomicrobium orale*^28^*, Desulfovibrio fairfieldensis*^29^ *and Treponema denticola*^25^. Conversely, decreased abundance of the genera *Moraxella* and *Capnocytophaga*^3^, as well as the bacterial species *Pasteurella multocida*^7^ is also commonly observed in periodontal disease in cats.

Our CLR-based compositional abundance analysis detected a multitude of microbes that are significantly upregulated or downregulated in feline periodontal disease. Our analysis confirmed the microbiome composition patterns associated with periodontal disease in published literature and also allowed us to identify additional microbes playing an important role in periodontal disease (**Table 1**).

**Table 1.** Selected microbial species which show significantly increased or decreased compositional abundance in periodontal disease compared to control (p<0.05). The average percentage increased or decreased abundance for each microbial species when compared to a healthy control (calculated using a centered log-ratio transformation) is shown in pink and blue, respectively. Microbial species previously described in scientific literature as misregulated in periodontal disease are shown in bold font.

| Microbes with increased abundance in PD cohort | % increase compared to Healthy cohort | Microbes with decreased abundance in PD cohort | % decrease compared to Healthy cohort |
| --- | --- | --- | --- |
| Bacteroides sp. HF-5287 | +49% | Frederiksenia canicola | -47% |
| **Bacteroides zoogloeoformans** | +47% | **Moraxella bovis** | -33% |
| Bacteroides sp. M10 | +41% | Mannheimia haemolytica | -32% |
| Odoribacter splanchnicus | +38% | Pseudoleptotrichia goodfellowii | -32% |
| Desulfobulbus oralis | +36% | Streptobacillus moniliformis | -32% |
| Bacteroides caccae | +35% | **Capnocytophaga sp. H4358** | -29% |
| **Desulfomicrobium orale** | +35% | **Capnocytophaga sp. H2931** | -29% |
| Bacteroides sp. CBA7301 | +33% | **Moraxella catarrhalis** | -28% |
| Bacteroides uniformis | +33% | Alysiella filiformis | -28% |
| Parabacteroides distasonis | +33% | **Moraxella cuniculi** | -27% |
| Bacteroides ovatus | +32% | **Moraxella ovis** | -27% |
| Bacteroides caecimuris | +32% | **Moraxella bovoculi** | -27% |
| **Desulfovibrio fairfieldensis** | +26% | Neisseria zoodegmatis | -27% |
| **Porphyromonas gingivalis** | +26% | Neisseria weaveri | -26% |
| Bacteroides heparinolyticus | +25% | **Capnocytophaga cynodegmi** | -25% |
| Actinomyces sp. Chiba101 | +25% | Neisseria animaloris | -25% |
| Bacteroides thetaiotaomicron | +25% | Cutibacterium acnes | -24% |
| Paraprevotella xylaniphila | +25% | Neisseria chenwenguii | -23% |
| Actinomyces howellii | +25% | Neisseria elongata | -22% |
| Bacteroides xylanisolvens | +24% | Neisseria dentiae | -22% |
| Bacteroides helcogenes | +24% | Kingella oralis | -22% |
| Petrimonas mucosa | +24% | Neisseria canis | -22% |
| Desulfovibrio desulfuricans | +24% | Pelistega sp. NLN63 | -21% |
| Bacteroides fragilis | +23% | Neisseria wadsworthii | -21% |
| Bacteroides sp. A1C1 | +22% | **Moraxella osloensis** | -21% |
| Treponema sp. OMZ 838 | +22% | **Capnocytophaga canimorsus** | -21% |
| Proteiniphilum saccharofermentans | +22% | Epilithonimonas vandammei | -19% |
| Treponema brennaborensense | +22% | Lysobacter oculi | -19% |
| Treponema putidum | +22% | Streptococcus dysgalactiae | -18% |
| **Treponema denticola** | +21% | Riemerella anatipestifer | -18% |
| Treponema pedis | +11% | **Capnocytophaga stomatis** | -17% |
| Acidovorax monticola | +11% | Fusobacterium hwasookii | -17% |
| Propionibacterium freudenreichii | +11% | Cardiobacterium hominis | -17% |
| Treponema phagedenis | +11% | Acinetobacter johnsonii | -17% |
| Prevotella denticola | +10% | Neisseria shayegani | -16% |
| Acidovorax sp. RAC01 | +10% | Fusobacterium<br>pseudoperiodonticum | -16% |
| **Tannerella forsythia** | +10% | **Pasteurella multocida** | -16% |

### Identification of dental disease predictive microbes whose compositional abundance is associated with periodontal disease, tooth resorption and halitosis

Having replicated existing findings describing the oral microbiome signature of periodontal disease, we then wanted to investigate if we could achieve a reliable population-based separation between the oral microbiomes of healthy cats and those suffering from periodontal disease, tooth resorption or halitosis. We identified 108 predictive microbes for periodontal disease, 74 for tooth resorption and 182 for halitosis, based on PLR microbial abundance comparisons between healthy/control oral microbiomes and those of cats suffering from one of the three dental conditions (**Supplementary table 1**). Plotting the average log ratio difference between significant pairwise microbial interactions in a dental condition versus control samples allowed us to separate sample populations based on their dental disease status (**Figure 2**). However, we also observed some overlap between the populations, meaning that for a certain set of samples, their compositional abundance of predictive microbes could be interpreted as either consistent with the control population or the respective dental disease population.

**Figure 2.**
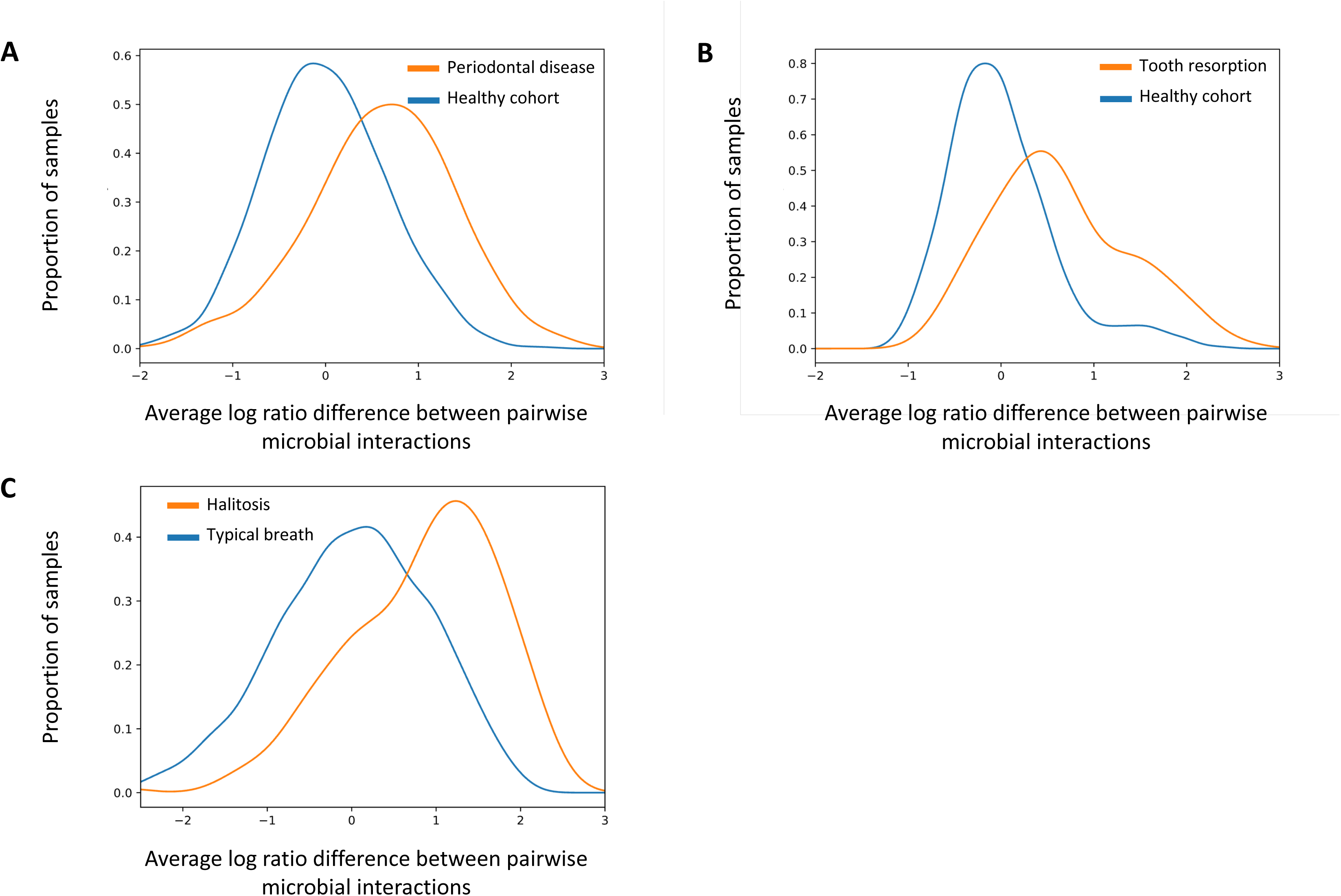
Distribution of the average log ratio difference scores between pairwise microbial interactions associated with (A) periodontal disease (PD) and healthy cohorts, (B) tooth resorption (TR) and healthy cohorts, and (C) bad breath (BB) and typical breath (TB) cohorts.

Next, we used 2 component Gaussian mixture modeling and plotted the probability that each dental disease and control sample is classified as belonging to one of the dental condition categories or to a control category based on each sample’s compositional abundance of predictive microbes (**Figure 3A, 3B, 3C**). We observed a bimodal probability distribution consistent with sample identity between dental condition and control for periodontal disease and halitosis and a weaker bimodal pattern for tooth resorption versus control. In all three instances, there was a minority of disease samples forming a small peak closer to 0 and a small set of control samples forming a slight peak closer to 1. This suggests that it is possible that a small proportion of the cats in our dental disease cohorts might actually be healthy or in remission (due to old, wrong or incomplete health information provided by the pet owner), while some cats in our control cohorts could be suffering from a dental condition that has not yet been diagnosed or noticed. We tested the sensitivity (ability to detect cats known to suffer from a dental condition) and specificity (ability to detect cats in the control cohort as not suffering from a dental condition) of our risk classification method for each dental condition (**Figure 3D**). Our method’s sensitivity is highest for halitosis and lowest for tooth resorption, while the specificity is highest for tooth resorption and lowest for halitosis.

**Figure 3.**
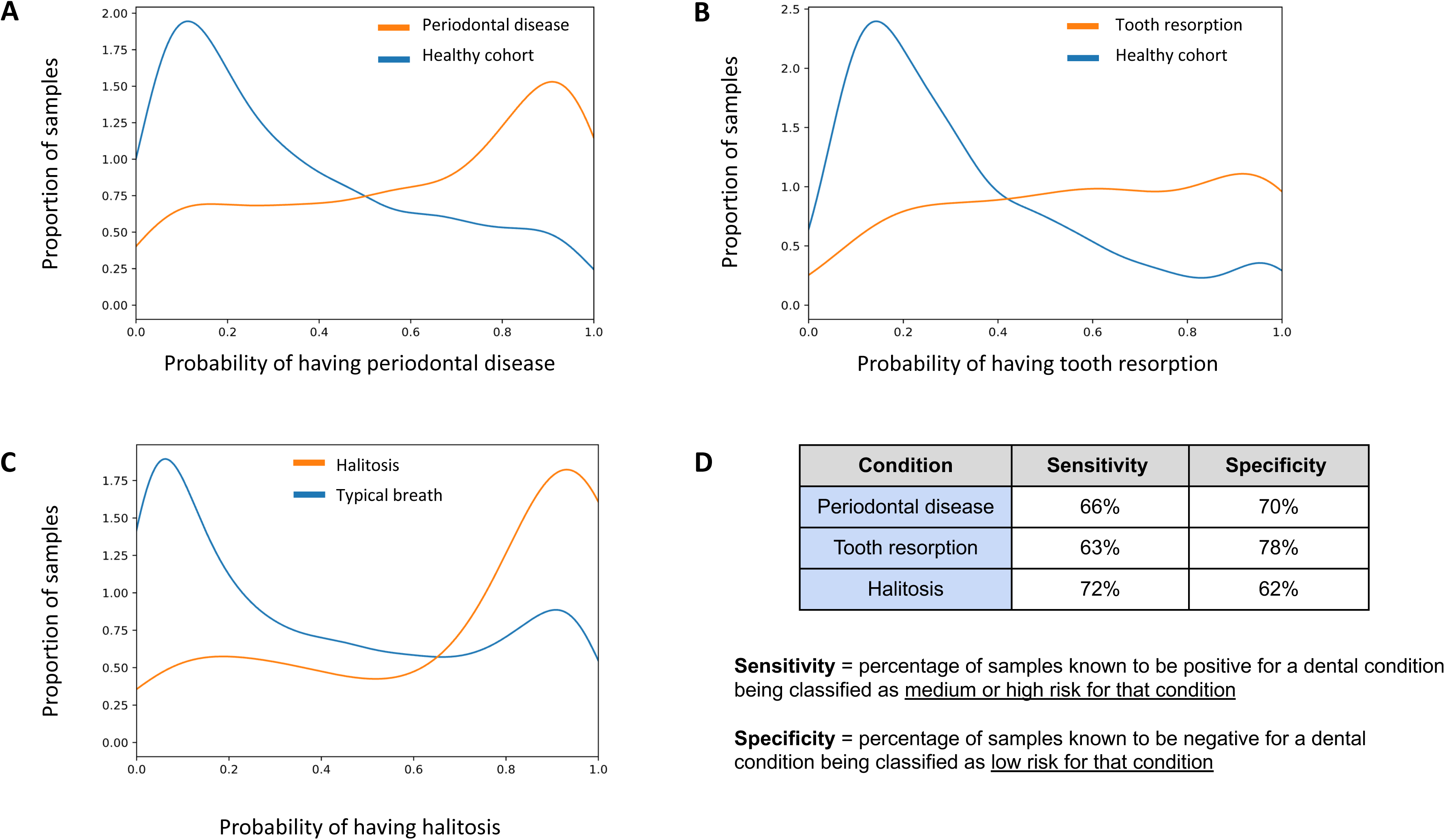
Sensitivity and specificity of the feline dental health test based on a 2-component Gaussian mixture model. Distribution of the probability of (A) cats from the PD and healthy cohorts being classified as having periodontal disease or being healthy; (B) cats from the TR and healthy cohorts being classified as having tooth resorption or being healthy; (C) cats from the BB and TB cohorts being classified as having bad breath or typical breath according to a 2-component Gaussian mixture model. (D) Sensitivity and specificity of the feline dental health test based on the ability to detect oral microbiome signatures characteristic for periodontal disease, tooth resorption and halitosis.

Our analysis identified 27 microbial species whose compositional abundance is predictive of all three dental conditions (**Figure 4**). While there was some overlap in predictive microbes between conditions, we also observed that each condition had its own specific set of predictive microbes, differentiating it from other conditions.

**Figure 4.**
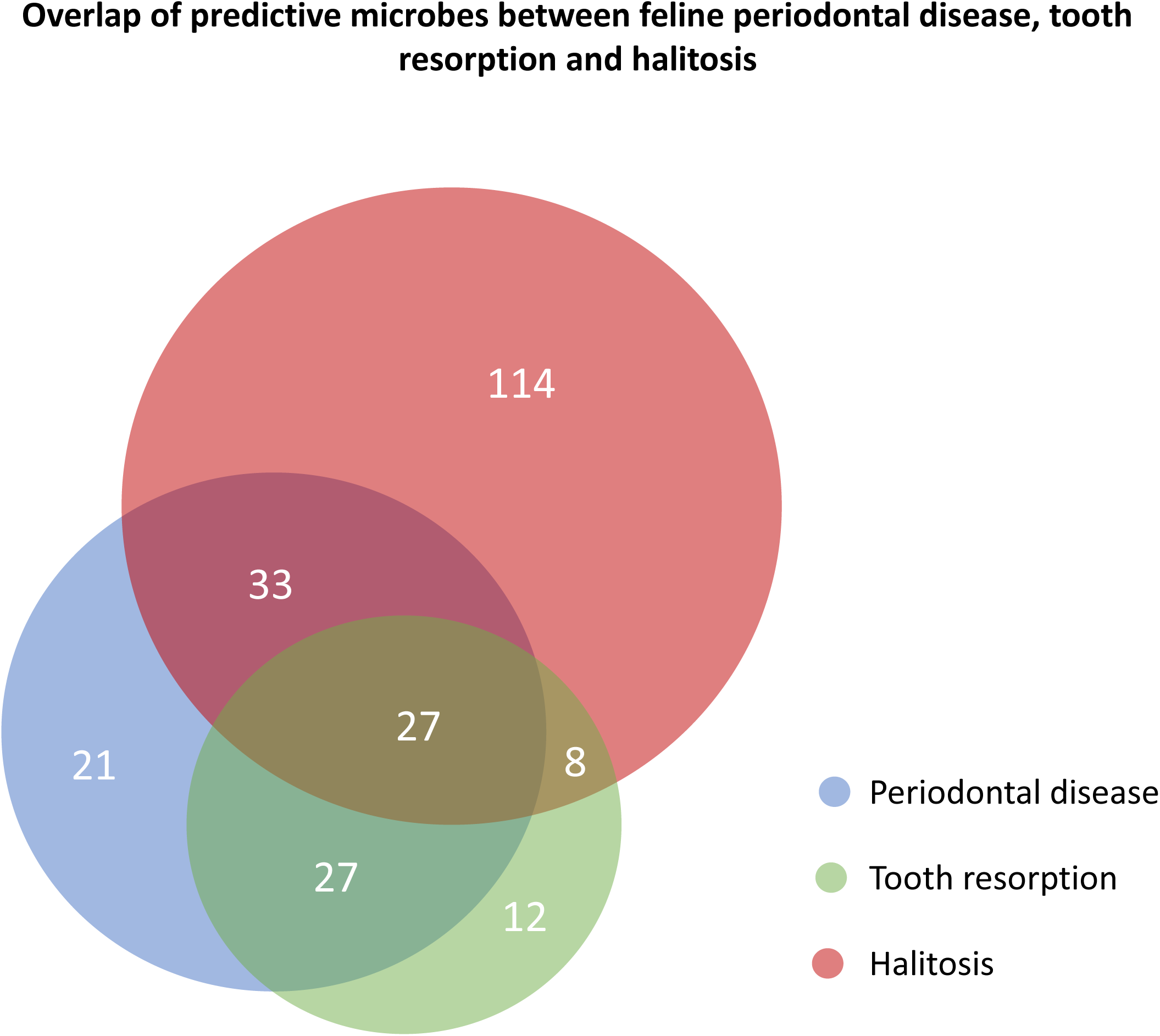
Overlap of oral microbiome predictive microbes characteristic of feline periodontal disease, tooth resorption and halitosis.

### Sampling location importance and consistency of the dental health test’s results

In order to investigate the reproducibility of our test’s results, we conducted two studies with volunteering cat owners. Study 1 asked pet owners to collect two oral swab samples from their cat - one targeting the whole mouth and the other - targeting specifically the top and bottom gum lines. Study 2 asked pet owners to collect three oral swab samples from their cat, each one targeting the whole mouth for sample collection. Study 1 had 12 participating cats and cat owners, while Study 2 had 11. There was a cat in Study 1 (Cat #12) that was on a daily doxycycline and prednisolone treatment following a full mouth extraction due to stomatitis. Due to the effect of the antibiotic treatment on the oral microbiome, we were not able to obtain a sufficient amount of microbial reads to generate a dental health report for Cat #12. For the remaining 11 cats in Study 1, using Spearman’s rank correlation, the ‘whole mouth’ and ‘gum line’ samples from the same cat clustered together in only two out of eleven cases, indicating that there is substantial variability between ‘whole mouth’ versus ‘gum line’ sampling methods (**Figure 5A**). However, the risk assessment for periodontal disease, tooth resorption and halitosis for the same cat mostly agreed between sampling methods. Wherever a discrepancy was observed, it was either medium risk in ‘gum line’ and low risk in ‘whole mouth’ (Cat #5, Cat #8, Cat #10), or high risk in ‘gum line’ and medium risk in ‘whole mouth’ (Cat #4). We did not observe strong discrepancies where one collection method indicated low risk for a dental condition, while the other collection method indicated high risk.

**Figure 5.**
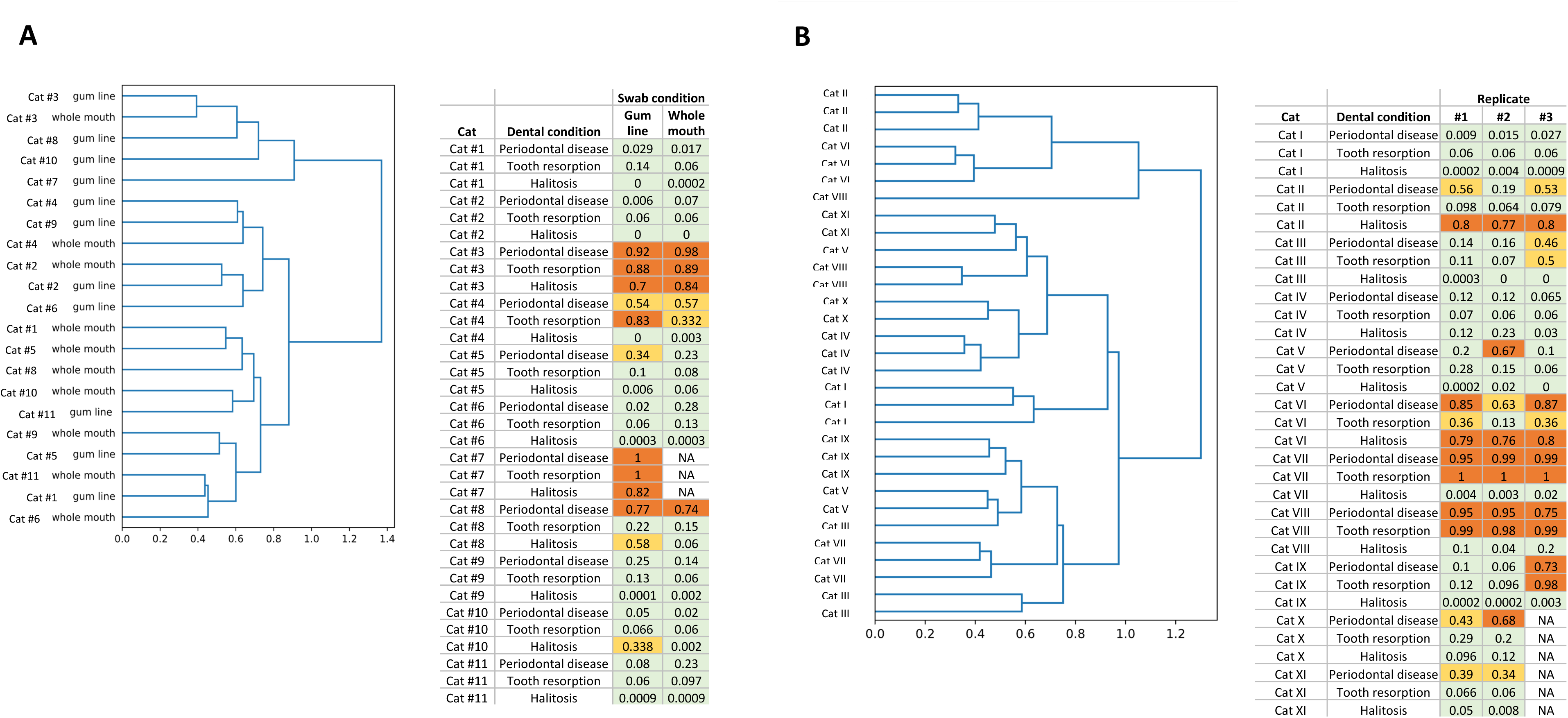
Sampling location effect and reproducibility of the feline dental health test results. (A) Results from Study 1 comparing the oral microbiome profiles of 11 cats based on sample collection methods targeting the whole mouth area or the gum line specifically. The dendrogram shows sample clustering based on Spearman’s rank correlation of the oral microbiome profiles. The table shows each participating cat’s risk assessment for periodontal disease, tooth resorption and halitosis based on the swabbing condition. Greed color indicates low risk, light orange - medium risk and dark orange - high risk. Cat #7’s ‘whole mouth’ sample was excluded from the analysis because the number of classified microbial reads was <10,000. (B) Results from Study 2 comparing the oral microbiome profiles of 11 cats based on three separate sample collections targeting the whole mouth. The dendrogram shows sample clustering based on Spearman’s rank correlation of the oral microbiome profiles. The table shows each participating cat’s risk assessment for periodontal disease, tooth resorption and halitosis based on each replicate. We received only two samples for Cat X from her pet owner. One of Cat XI’s samples was excluded from our analysis because the number of classified microbial reads was <10,000.

In contrast, Spearman’s rank correlation analysis indicated that all three ‘whole mouth’ triplicates from the same cat clustered together in 8 out of 11 cases (**Figure 5B**). However, unlike the results from Study 1, we observed two cases of more noticeable discrepancies between triplicates. One of Cat IX’s samples indicated that the cat was at high risk for periodontal disease and tooth resorption, while results from the remaining two samples suggested she was at low risk for those two conditions. In addition, one of Cat V’s sample results indicated he was at high risk for periodontal disease, while his remaining two samples indicated he was at low risk for this condition. In Study 2, where participants were instructed to collect a ‘whole mouth’ swab sample, uneven sampling of areas where most dental diseases are manifested (the gum line) could potentially explain the discrepancy between some of the triplicates.

## Discussion

We developed a painless and easy to use oral swab based feline dental health assessment test that can be administered either at home by the pet owner or at the veterinary clinic or office. Our dental health test can serve as a powerful tool for pet owners to monitor their cat’s dental health and detect any disease-related oral microbiome changes early on. Using this test for routine screening could help reduce the number of emergency feline dental visits, facilitate diagnosis of dental conditions earlier (when more treatment options available), and decrease the prospect of cats experiencing severe dental pain as a consequence of advanced dental disease going undiagnosed for long periods of time.

Our approach used shotgun metagenomic sequencing for the identification of microbes associated with dental disease in cats. Since available literature on the oral microbiome characteristics of cats suffering from tooth resorption or halitosis is limited, we focused on the microbial signature of periodontal disease to validate our method. We observed a significantly increased compositional abundance of *P. gingivalis, T. forsythia, B. zoogleoformans, D. orale, D. fairfieldensis* and *T. denticola* (among other microbes) in the microbiomes of cats suffering from periodontal disease. Furthermore, we also observed significantly decreased compositional abundance of the genera *Moraxella* and *Capnocytophaga*, as well as the bacterial species *P. multocida*. These observations are all consistent with previous findings from studies focused on the oral microbiome of cats, humans and dogs suffering from periodontal disease ^3, 4, 7–9, 24–29^. These results lend strong support to the use of shotgun metagenomic NGS in combination with compositional abundance analysis for the identification of microbes associated with different dental diseases in cats. Our feline dental health test relies on a comparison of the cat’s current oral microbiome state to the oral microbiomes of cats reported by their pet owners to have been diagnosed with periodontal disease or tooth resorption or to suffer from bad breath. The comparison is based on the compositional abundance of microbes determined by our analysis to be predictive of each of the three dental conditions. It is important to note that the use of the word ‘predictive’ is not meant to be interpreted as ‘causative’, it simply reflects the fact that a microbe has a significantly different compositional abundance in a particular dental condition compared to control. This could either mean that the microbe has an active role in the disease’s pathology or that the changes of its compositional abundance are a byproduct of pathology. This distinction is particularly important to stress in the case of tooth resorption, where there is currently no peer-reviewed literature that has identified a microbial cause for this disease.

In veterinary practice, dental disease is sometimes thought of as a syndrome where halitosis, tooth resorption and periodontal disease are rarely seen separately from each other, even though they can have different underlying pathologies. This view is, to some extent, reflected in our data where we see some overlap in these microbes between conditions. The largest overlap is between halitosis and periodontal disease, which is consistent with observations from the clinic where halitosis is often a harbinger of periodontal disease^30, 31^. However, we also identified a plethora of microbes whose compositional abundance in the oral microbiome is predictive specifically of halitosis, tooth resorption or periodontal disease. This suggests that there are microbial profiles associated with specific dental pathologies, in addition to the existence of a core set of microbes associated with dental disease in general.

Our test has different sensitivity and specificity for each of the three conditions, with the highest sensitivity for halitosis (72%) and the highest specificity for tooth resorption (78%). The model’s sensitivity is lowest for tooth resorption (63%). This could be due to the nature of the pathology behind tooth resorption. It tends to originate inside of the tooth and, as it enters more advanced stages, it then reaches the surface of the tooth^32^. It is possible that the microbes associated with tooth resorption can be detected most reliably when the resorptive process has reached the surface of the tooth. This could explain why our model might potentially miss some earlier stage cases of tooth resorption.

Our model’s specificity for periodontal disease and bad breath is lower (70% and 62%, respectively) compared to the test’s specificity for detecting tooth resorption associated changes in the microbiome (78%). This observation could be explained by the possibility that our healthy and TB cohorts include some cats with periodontal disease or bad breath, respectively, that have not yet been diagnosed by a veterinarian or noticed by the pet owner. However, even with these caveats in mind, the specificity and sensitivity of our feline dental health test for all three conditions are comparable to (or better than) previously reported human microbiome-based disease risk assessment algorithms^15, 33^.

Our method’s specificity and sensitivity is potentially also influenced by the sample collection method. Our current risk prediction models are based on pet owner-provided oral swab samples where the whole mouth was targeted for sample collection, focusing on no particular area of interest. As our replicate studies have shown, risk assessments based on a ‘whole mouth’ swab sample can occasionally show variability (specific examples are Cat V’s and Cat IX’s samples). This is probably due to the fact that when the pet owner is instructed to collect a ‘whole mouth’ swab sample, different mouth areas get preferentially swabbed each time. Since the easiest area to swab is the tongue, it is possible that some ‘whole mouth’ sample collection attempts focus on the tongue area where the presence of dental disease-associated microbes is more variable compared to the gum line. Interestingly, when there was a discrepancy between ‘whole mouth’ and ‘gum line’ targeted oral swab collection in our Study 1, the ‘gum line’ risk score was always higher than the ‘whole mouth’ risk score (Cat #4, Cat #5, Cat #8, Cat #10). This supports our hypothesis that sample collection targeted at the gum line is likely to more accurately represent microbiome states linked to dental diseases. For this reason, to maximize chances of identifying dental disease-associated microbiome signatures, we are strongly advising every pet parent and veterinarian that uses this dental health test to collect an oral swab across the cat’s gum line instead of their whole mouth.

Some of the limitations of our work are focused around the fact that, even though we used a sizable domestic cat cohort (n=6,110), the health history data for these cats was provided by the pet owner. Despite the fact that we asked pet owners if their cats had been diagnosed by a veterinarian with periodontal disease or tooth resorption, some of the diagnostic precision would have, undoubtedly, suffered, having been relayed by the pet owner. To alleviate this problem and limit instances where a cat reported by their pet owner to be healthy had actually started developing a yet undiagnosed dental disease, we set an age limit to our control healthy cohort of 1-3 years. This limit was set due to the well-established connection between age and dental disease^31, 34^. A potential drawback of this age restriction could be that our healthy control cohort could be biased towards the oral microbiomes of younger cats and not be representative of older cats with no dental or systemic diseases. Lastly, the assessment of whether cats in our BB and TB cohorts had halitosis or not was based on the subjective evaluation of the pet owner, which could have potentially added another source of bias.

We are routinely updating our database and including more cats suffering from periodontal disease, tooth resorption and halitosis for the continuous optimization of our test’s specificity, sensitivity and reproducibility. Our next step is to gather oral microbiome data directly from feline patients at the veterinary clinic, where their most current and precise diagnoses can be obtained. This will lead to the inclusion of other dental diseases to this test, for example gingivostomatitis, and further improving our model’s sensitivity, specificity and reproducibility.

An exciting future direction for the study of the feline microbiome could be performing a *de novo* metagenomic assembly on our shotgun sequencing data. This could allow us to discover microbial species not previously described in existing databases. Some of these species could potentially have significant associations with dental and systemic health conditions. Additionally, the metabolic output of the oral microbiome could also be simulated using known enzymatic pathway analysis tools. This analysis would provide an additional dimension to the existing microbial composition data to further characterize disease signatures and improve predictive disease models.

## Acknowledgements

We would like to thank Jan Bellows, DVM, DIPACVD (canine and feline specialty), DIPABVP for our discussions on feline dental health and his comments on this manuscript. We would also like to thank Basepaws customers who agreed for us to use their cat’s data for research purposes.

---

**Supplementary figure 1. Microbial species richness as a function of number of sequencing reads - ligation-based NGS library preparation versus tagmentation-based NGS library preparation method.**

**Supplementary table 1. Predictive microbes for periodontal disease, tooth resorption and halitosis based on pairwise log-ratio microbial abundance comparisons between healthy/control oral microbiomes and those of cats suffering from one of the three dental conditions.** ‘1’ indicates that the microbes is considered predictive of a particular dental condition, while ‘0’ indicates that it is not.

