## Supplementary figures and images for "Development of an oral swab based microbiome test for the detection of feline dental disease"

### Supplementary figure 1

**Supplementary figure 1**

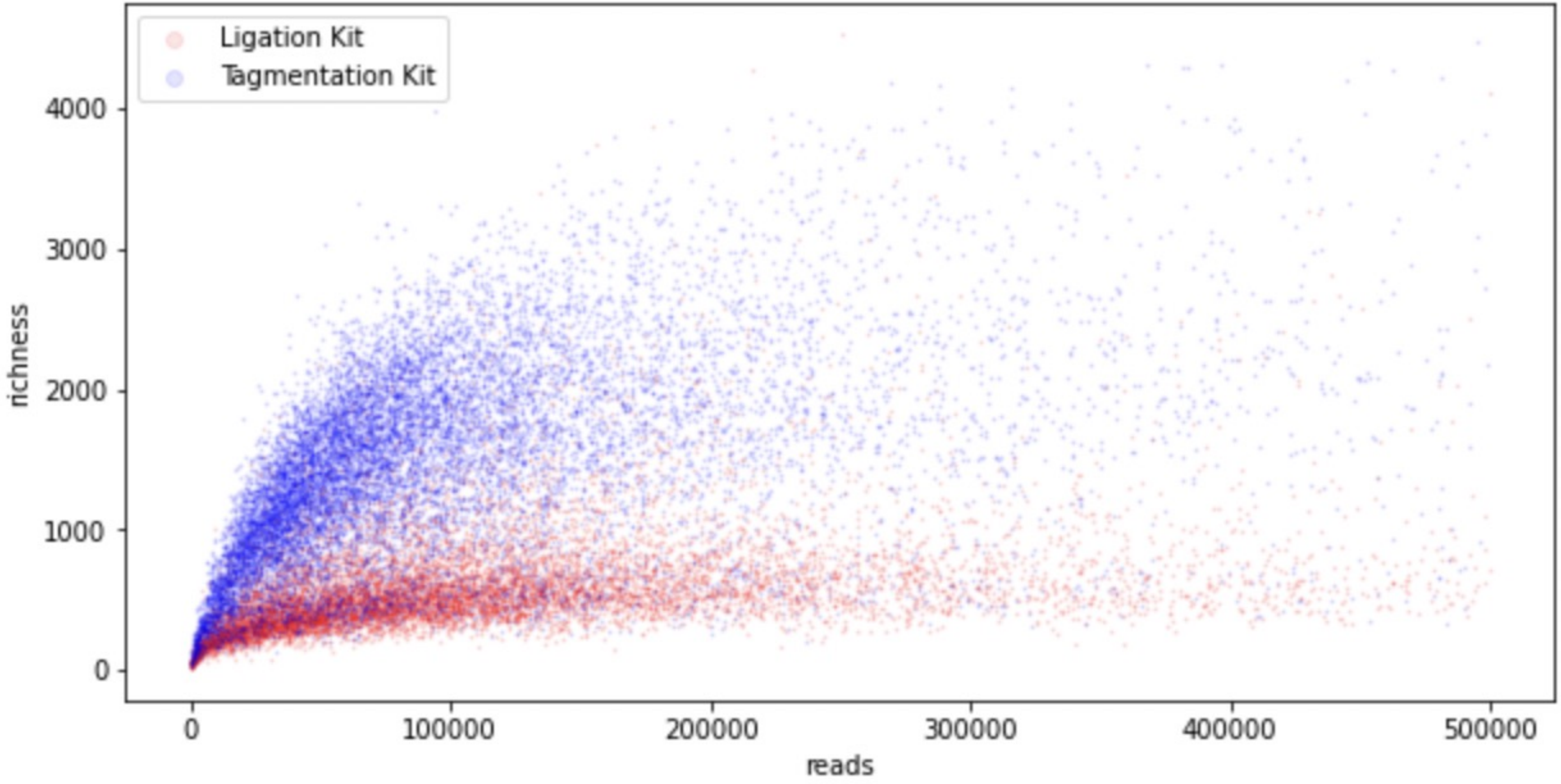
